# Developing methods for measuring national distributions and densities of wild mammals using camera traps: A Kosovo study

**DOI:** 10.1101/2020.07.30.193078

**Authors:** Sarah E. Beatham, Alastair I. Ward, David Fouracre, Jeton Muhaxhiri, Michael Sallmann, Besim Zogu, Valdet Gjinovci, Anthony J. Wilsmore, Graham C. Smith

## Abstract

Understanding the distributions and density of wild mammals is integral to the implementation of wildlife management strategies, particularly for controlling diseases and conservation management. Recent advances in camera trap technology together with the development of the Random Encounter Model have provided a non-invasive method for estimating mammal densities. In addition, the development of citizen science initiatives have advanced ecological data collection. This study describes a national camera trap survey delivered by local stakeholders in eleven forest sites in Kosovo from 2014 to 2015 to measure the distributions and abundance of medium to large wild mammals as part of the Control and/or eradication of animal diseases project. The Random Encounter Model was used to calculate density data for each species, which appear realistic when compared to densities found in other European countries. The study particularly focussed on the red fox (*Vulpes vulpes*) and the grey wolf (*Canis lupus*) as potential vectors of rabies and wild boar (*Sus scrofa*) as a vector of classical swine fever. These species were found to be three of the most widely distributed species in Kosovo and were present at the majority of sites at high densities. The camera survey also provided information on species of conservation concern such as the Eurasian brown bear (*Ursus arctos*) and provided the first physical evidence of a live Eurasian golden jackal (*Canis aureus*) in Kosovo. Although sources of bias were identified, these estimates are likely to be more accurate than those devised from methods such as hunting bags and the findings of this study suggest that, with a moderate amount of development, camera trapping implemented by local stakeholders can be used as an effective and practicable method to estimate national distributions and population sizes of medium to large sized wild mammals.

## Introduction

Understanding the distributions and density of wild mammals is integral to the design and implementation of wildlife management strategies for disease control, hunting and conservation management. This information can be used to provide guidance on selecting the most appropriate strategy to use, the geographical scale over which to use it and the effort required to meet the desired objectives.

One of the largest and most successful wildlife disease control campaigns has been the eradication of rabies from much of Europe through successive oral vaccination campaigns implemented from the 1970s onwards. The red fox (*Vulpes vulpes*) has the widest geographical distribution of any member of the family Carnivora [1] and it is also the main vector for wildlife rabies in Europe, contributing to more than 80% of cases [2]. Since 2008, efforts have been made to control and eradicate rabies and classical swine fever (CSF), a disease of wild boar (*Sus scrofa*), in the Western Balkans, a region in South East Europe which borders European Union (EU) states and where rabies is endemic [3]. There is now also concern over the distribution and abundance of wild boar across Europe for the risk assessment relating to African Swine Fever [4] and there is a paucity of data in Eastern Europe [5].

Kosovo is the smallest of the Balkan countries, declaring itself independent from Serbia in 2008. There has been limited rabies surveillance in the country, although two cases were recorded in foxes in 2007 near to the border of the Former Yugoslav Republic of Macedonia (FYROM) [6]. The control and/or eradication of animal diseases (DCE) project was established in Kosovo in 2010, funded by the EU and managed by the EU Office in Kosovo, with the Kosovo Food and Veterinary Agency as its main beneficiary. Two of the project’s objectives were the monitoring of an oral rabies vaccination campaign and the surveillance of wildlife for rabies and CSF. To help meet these objectives, up to date data on the distribution and densities of wild mammals in Kosovo were obtained, focussing the red fox, grey wolf (*Canis lupus*) and wild boar. The grey wolf may also be a rabies vector and a rabid wolf was reported in Macedonia in 2011 [7]. The wolf is not regarded as a reservoir host, but rabid wolves may travel long distances and inflict serious injuries to humans and other animals, facilitating disease transmission [8].

There is currently limited information on wild mammals in Kosovo. The last population estimates, recorded in 2003, were derived from observations from land management experts (e.g. hunters) and focussed on species of economic and/or conservation importance [9]. Similarly, there are no European countries that have up to date, accurate and comprehensive data on the distributions and densities of wild mammals, although Great Britain has recently published a ‘systematic approach to estimate the distribution and total abundance of British mammals’ [10]. Information on the national distributions of wild mammals often comes from data on hunting bags or direct counts by hunters which, while having the benefits of being relatively low cost, low effort and non-invasive, are subject to a number of biases [11]. Recent advances in camera trap technology together with the development of the Random Encounter Model (REM) have provided a non-invasive method for estimating mammal densities [12]. In addition, the development of citizen science initiatives has enabled the use of more resources for the collection of increased quantities of ecological data when compared to other approaches [13]. This study describes the findings of a national camera trap survey implemented in Kosovo forests from 2014 to 2015 by local stakeholder groups, to measure the distributions and abundance of medium to large wild mammals. The camera method is evaluated based on the apparent validity of the results obtained and practicability compared with existing methods used to measure mammal distributions and densities and suggestions are made on how the method could be further improved.

## Methods

### Study area

Kosovo has a human population of 1.8 million, an area of 10,908 km^2^ and lies between 41° and 44° N and between 20° and 22° E and 270 m to 2656 m above sea level. Its borders are predominantly mountainous and neighbours include Serbia to the North and East, Montenegro to the North West, Albania to the South West and FYROM to the South. Covering an area of 4,810 km^2^, forest is Kosovo’s largest land cover type [14]. Eleven forest sites were selected as representative of Kosovo forest in terms of topography, flora, geographical range, altitude and management (Fig 1). Included were the private licensed hunting grounds of Shtime, Pristina, Malisheva and Kamenica, the municipal authority owned areas of Jezerc, Podujevo and Mitrovica, the Kosovo Forest Agency owned site of Dubocak, the Kosovo Ministry of Environment and Spatial planning owned “Bjeshkët e Nemuna” National Park areas of Istog and Junik and the Sharri national park area of Dragash. The areas sampled primarily consisted of deciduous forest, though small areas of coniferous species and grassland at forest edges were also included. In Dubocak, Istog, Junik and Dragash, hunting was permanently prohibited. Hunting was prohibited at all of the study sites from January 2015 when a national ban was introduced by the Kosovan Government in an attempt to control illegal hunting. This remained in place for the duration of the study.

**Fig 1.**
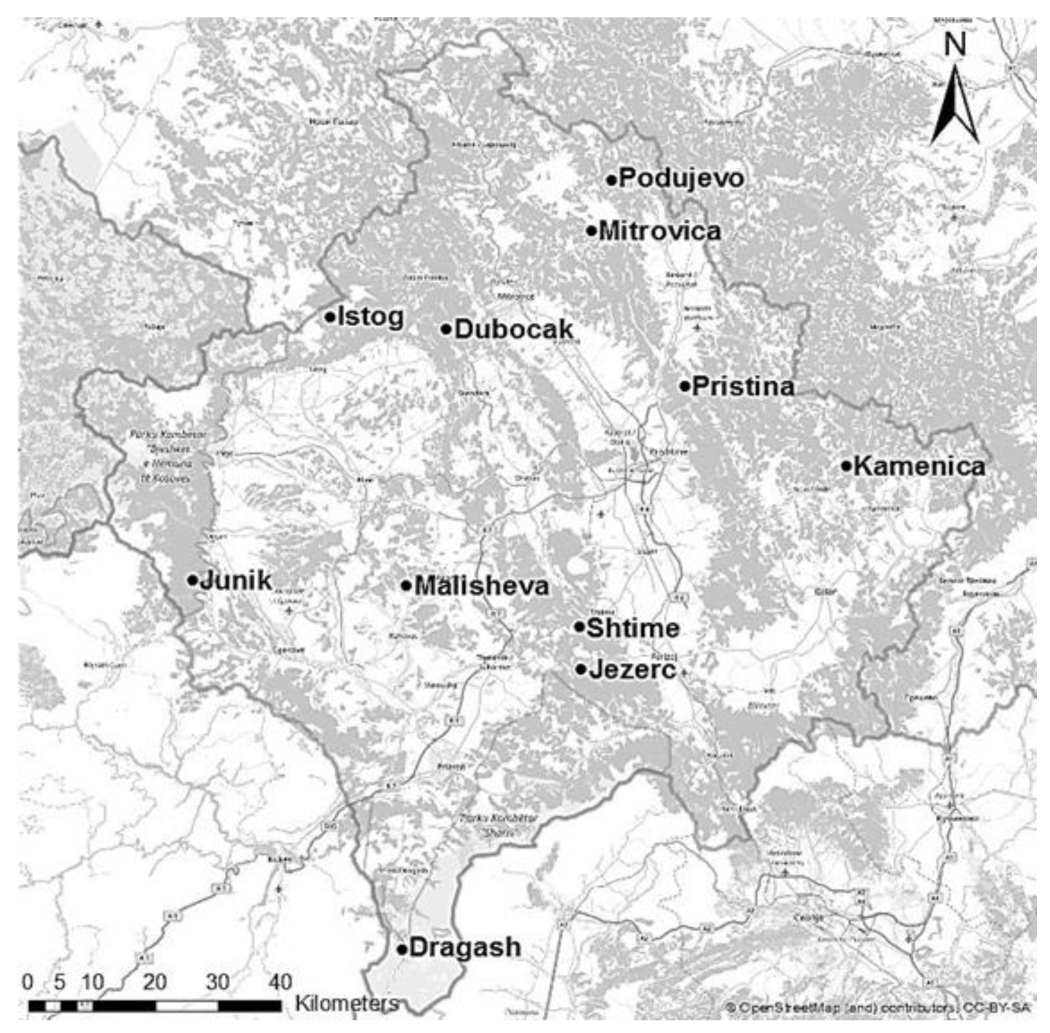
Eleven study sites in Kosovo at which cameras were deployed and land manager surveys performed. Forest areas are shaded grey. Background map from ©OpenStreetMap contributors, data available under Open Database Licence: www.openstreetmap.org/copyright.

### Camera deployment and mammal identification

Camera trap deployment was led by the DCE project team in accordance with written guidelines and training on how to deploy a transect of cameras in a forest environment provided by the National Wildlife Management Centre, APHA, UK. Cameras were deployed by a different stakeholder group at each site, which included the Kosovo Forestry Agency, local hunting associations and US members of the Kosovo Force (KFOR). Stakeholder groups provided resources including man hours, 4×4 vehicles and access to each forest area. On each occasion camera deployment was supervised by representatives from the DCE project team.

The camera deployment method was designed to obtain a representative sample of mammal records for each area of forest given limited available resources in terms of number of cameras and man hours available. From July 2014 to February 2015, ten Moultrie^®^ Game Spy D55IR (EBSCO Industries Inc., USA) (D55IR) cameras were deployed over one or two transects across each area of forest, for between 11 and 15 days at each site. To increase comparability, the same camera units were used for each site. Camera location was guided by level of accessibility, perceived risk of theft and availability of suitable places to attach cameras. A Garmin eTrex® 10 GPS unit (Garmin Ltd, UK) was used to record location coordinates for each camera. Cameras were secured to trees between 0.5 and 1.0 metres above the ground and were placed to maximise the chance of capturing any medium to large wild mammals passing. The cameras were set so that, when they were triggered through motion or heat, they took one photograph and one video. The video length was set at between 5 and 15 seconds. The delay between successive triggers was set at 5 minutes to minimise the occurrence of multiple triggers from the same animal or group of animals.

From March 2015 to December 2015, the deployment methodology was modified to increase data collection with 19 or 20 cameras deployed for 20 to 24 days and an additional camera, the Moultrie^®^ M-880 Gen2 (EBSCO Industries Inc., USA) (M-880) was used. The M-880 had improved battery performance, permitting video for up to one minute. Camera placement was standardised through deployment at a distance of approximately one camera every 330 m. Two transects were deployed across two separate locations within each study site to ensure a more random and representative sampling.

All photographs and videos were analysed from each camera and all medium to large (average body weight >1 kg) sized wild mammals recorded. Species identification was assigned only after an agreement was reached by at least two wildlife experts. If a mammal could not be identified from its photograph, the associated video was used to aid identification.

### Estimation of mammal density

The population density (D) measured as individuals/km^2^ for each species at each study site was calculated using equation 4 of the Random Encounter Model [12]:

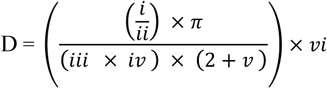

Each component of the equation was calculated as follows:

i)Number of records for each species: each photo and its associated video constituted a single record for that mammal, even if more than one individual was present. Two species of *Martes*, pine marten (*Martes martes*) and beech marten (*Martes foina*), were often indistinguishable, so were treated as a single species.

ii)Number of camera days: the sum of the number of days each camera was deployed, taking account of any faulty or stolen cameras.

iii)Speed of movement (km/day): the mean daily range in km/day taken from radio-telemetry or GPS data from a published study or existing dataset, assessed as the most representative in terms of location, environment and individuals sampled.

iv)Radius of detection (km): the maximum distance over which a species could be detected and identified. This was measured empirically for each model of camera by a walk-test performed by a person at increasing distances in front of the camera. As 66% of the species records were obtained at night, the walk-test to determine the radius of detection was executed in a room without any light source. In addition, all of the photographs that contained clearly identifiable mammals were analysed to visually estimate the distance from the camera, to the closest metre, at which detection and identification was possible for individuals from each species. Only photographs containing single individuals were used to ensure there was no ambiguity as to which individual triggered the camera. Additional information on factors influencing detection were recorded from the analysis of each photograph, including whether a forest track was present and a visual estimation of vegetation cover as a percentage of the field of view (to the nearest 25%).

v)Angle of detection (radians): the optical field of view, stated by the manufacturer, for each camera model. This was considered to be the maximum possible angle over which species could be both detected and identified.

vi)Average group size: the average group size for each species calculated from the camera videos. Only adults and juveniles approaching full development were included in the group size to reduce seasonal bias. Where video information was too limited, the average group size was taken from appropriate literature in terms of geography, methodology and individuals sampled.

Mean density estimates were calculated for each species using the REM and a 95^th^ percentile range for predicted values was calculated by bootstrapping the number of records associated with each camera location 10,000 times, using Crystal Ball (Oracle® Fusion Middleware, 2016). To provide a mean density estimate for each mammal species for forests in Kosovo, data from all study sites were combined and population size estimated by multiplying the density by forest area.

### Land manager surveys

Prior to this study, information on the distributions and relative abundance of wild mammals in Kosovo had been collated from assessments by the hunting community. To compare this more traditional approach with the camera survey method, additional information on the distribution and frequency of mammals was obtained through surveys conducted with private hunters, hunting associations or other land managers who oversaw the areas where the cameras were deployed. Land managers from each area were asked two questions; which medium to large sized wild mammals were present in their area and whether the species were frequent (twelve or more sightings per year of individuals or groups) or sporadic (one to eleven sightings per year of individuals or groups). The findings were compared with the results from the camera survey.

## Results

### Camera deployment

The details of the camera deployment at each study site are shown in Table 1. Twelve cameras were reported as stolen and therefore did not contribute any data, including six from Istog, one from Junik, four from Podujevo and one from Dubocak. Between July 2014 and February 2015, ten locations per site were sampled by cameras and the sum of the number of days the cameras were deployed for averaged at 137 days per site. From March 2015, after losses were accounted for, an average of 17 locations per site were sampled by cameras and the sum of the number of days the cameras were deployed for averaged at 350 days per site. This gave a total of 2,572 camera days across the 11 sites.

**Table 1.** Details of the camera deployment at Kosovo forest sites. Number of cameras and camera days take into account stolen or faulty cameras. Transect length is the total distance in km covered by the transect(s) of cameras at each site. Camera days is the sum of the total number of days each individual camera was deployed for.

| Site | Altitude (m) | | Transect length (km) | Number of working cameras | Transects | Distance between cameras (Mean ± SD) | Camera type | Camera days | Survey period |
| --- | --- | --- | --- | --- | --- | --- | --- | --- | --- |
|  | Min | Max |  |  |  |  |  |  |  |
| Malish. | 655 | 754 | 2.64 | 10 | 1 | 256 ± 75 | D55IR | 120 | Jul 2014 |
| Shtime | 730 | 912 | 4.37 | 10 | 1 | 388 ± 251 | D55IR | 140 | Aug 2014 |
| Jezerc | 736 | 1005 | 1.52 | 10 | 2 | 176 ± 51 | D55IR | 140 | Sep/Oct 2014 |
| Kamen. | 596 | 926 | 9.63 | 10 | 1 | 1201 ± 1019 | D55IR | 150 | Oct/Nov 2014 |
| Pristina | 655 | 858 | 14.6 | 10 | 1 | 1299 ± 666 | D55IR | 120 | Dec 2014 |
| Drag. | 928 | 1331 | 16.7 | 10 | 1 | 1522 ± 942 | D55IR | 150 | Jan/Feb 2015 |
| Duboc. | 749 | 992 | 6.27 | 19 | 2 | 311 ± 71 | D55IR | 414 | Mar to Jun 2015 |
| Junik | 1003 | 1352 | 5.16 | 18 | 2 | 249 ± 89 | M-880 | 378 | Aug 2015 |
| Istog | 1521 | 1897 | 5.57 | 13 | 2 | 297 ± 65 | M-880 | 260 | Sep/Oct 2015 |
| Mitrov. | 991 | 1609 | 7.09 | 19 | 2 | 311 ± 67 | D55IR /M-880 | 380 | Nov 2015 |
| Poduj. | 778 | 1191 | 4.05 | 16 | 2 | 204 ± 72 | D55IR /M-880 | 320 | Dec 2015 |

### Mammal identification and recording

In total, 376 photographs and associated videos contained an identified medium to large sized wild mammal. Additional photographs were obtained of people, domestic animals and rodents. For 20 photographs, it could not be determined what triggered the camera. In 7% of cases the video record was required in addition to the photograph to verify the species/genus. Overall, 97% (364) of mammal records could be identified to species level with reasonable certainty, or in the case of *Martes*, to genus level. Other species identified included roe deer (*Capreolus capreolus*), Eurasian wild boar, red fox, grey wolf, Eurasian badger (*Meles meles*), European hare (*Lepus europaeus*), European wildcat (*Felis sylvestris*), Eurasian brown bear (*Ursus arctos*) and the Eurasian golden jackal (*Canis aureus*). This was the first physical evidence of a live golden jackal in Kosovo.

The number of records obtained for each species from each site varied considerably (Supplementary Information S1 Table). There were 9 records per site for survey period 1 and 60 records per site for survey period 2. Over half the records from survey period 2 came from the site of Junik. The sites where the most species were recorded were Shtime and Junik, with eight species recorded at each and the sites with the lowest numbers of species recorded were Kamenica and Dragash, with two at each. The species recorded at the most sites were the red fox at nine sites, and the grey wolf and European hare at eight sites each. All species were recorded at a minimum of four different sites, with the exception of the golden jackal, which was recorded at Podujevo only.

### Estimation of mammal densities

Speed of movement values (Table 2) were obtained from a published study for seven species and from datasets from organisations sourced through the Movebank website, Euroungulates for roe deer and Biomove for European hare. The DDM for roe deer appeared low, therefore a second study was consulted. A radio-telemetry study [15] found the mean DDM for roe deer varied between 542 and 1856 metres, dependent on month and sex. The majority of individuals analysed had a DDM of less than 1 km, thus supporting the value of 800 metres used for the REM in this study.

**Table 2.** The sources used of the daily distance moved (DDM) including the mean and SD, the number of fixes per day, the number of individuals (N) including males (M) and females (F), method used, study period and location and habitat of the study.

| Species | DDM<br>(km day<sup>-1</sup>) |  | Fixes<br>day<sup>-1</sup> | N | Method | Study<br>Period | Location | Source |
| --- | --- | --- | --- | --- | --- | --- | --- | --- |
|  | Mean | SD |  |  |  |  |  |  |
| Red fox | 6.8 | 1.9 | 96 | 13<br>M = 3<br>F = 10 | Radio-telemetry | Sep 1989-Aug 1993 | Jura mountain<br>mixed habitat,<br>Switzerland | [16] |
| Grey wolf | 22.8 | 2.0 | 48 | 11<br>M = 2<br>F = 9 | Radio-telemetry | Mar 1994 -<br>Sep 1999 | Białowieża forest,<br>Poland | [17] |
| Wild boar | 6.8 | 2.6 | 48/96 | 20 | Radio-telemetry | Feb 2006-Dec 2008 | Białowieża forest,<br>Poland | [18] |
| Roe deer | 0.8 | 0.6 | 2 | 115<br>M = 61<br>F = 54 | GPS | Dec 2004 -<br>Apr 2012 | Bavaria forest,<br>Germany | www.euroun-<br>gulates.org |
| Eurasian<br>badger | 7.0 | 3.1 | 48/96 | 13 | Radio-telemetry | 1997-2000<br>(Excluding<br>winter) | Białowieża forest,<br>Poland | [19] |
| European<br>hare | 1.8 | 1.1 | 24 | 64<br>M = 33<br>F = 31 | GPS | May 2011-Jan 2016 | Uckermark and<br>Freising, mixed<br>arable, Germany | www.biomove-<br>.org |
| Pine<br>marten | 5.5 | N/A | 96 | 14<br>M = 6<br>F = 8 | Radio-telemetry | Apr 1991 -<br>Mar 1996 | Białowieża forest,<br>Poland | [20] |
| European<br>wildcat | 6.7 | N/A | 63 | 12<br>M = 6<br>F = 6 | Radio-telemetry | 1980 -1984 | Meine forest,<br>France | [21] |
| Eurasian<br>brown bear | 6.0 | N/A | 24 | 33<br>M = 19<br>F = 14 | GPS | 2005-2009 | Mixed habitat,<br>West Slovenia | [22] |

The majority of studies from which the values were sourced were located in central or eastern Europe and in forest habitats, including several located in the Białowieża forest, Poland. The estimates for each species were calculated from radio-telemetry data with the exception of brown bear, hare and roe deer where GPS data were used.

Using the walk-test, it was found that the maximum distance that would allow reliable detection of an individual was found to be eight metres for both camera types. The estimated average distance between the camera and all species photographed was 2.3 metres (SD 1.3) and the estimated maximum distance was between three metres (brown bear and wildcat) and eight metres (red fox). This was primarily influenced by the proximity of forest tracks to the cameras (apparent in 45% of the mammal records) and vegetation cover (which obstructed at least 25% of the field of view for 42% of the mammal records). Based on the data provided by both analyses, the radius of detection was set at eight metres.

The angle of detection was 0.907571 radians for the D55IR camera model and 0.872665 radians for the M-880 camera model. For the Kosovo forest species densities, when the ratio of D55IRs to M-880s was taking into account, the angle of detection was calculated proportionately as 0.892778 radians.

The average group size values were calculated from video records for eight species (Table 3). An average of 22 videos (minimum 6, SD 21) were used for each species from an average of 5.7 sites (minimum 3, SD 1.8). Group sizes ranged from one individual (fox, badger, marten and wildcat) up to 1.8 individuals for the grey wolf. For wild boar, it was found that individuals sometimes moved around in large groups which were often not entirely captured by the videos. Therefore group size for wild boar (four individuals) was taken from a study on wild boar social groups in the Camargue, southern France [23].

**Table 3.**
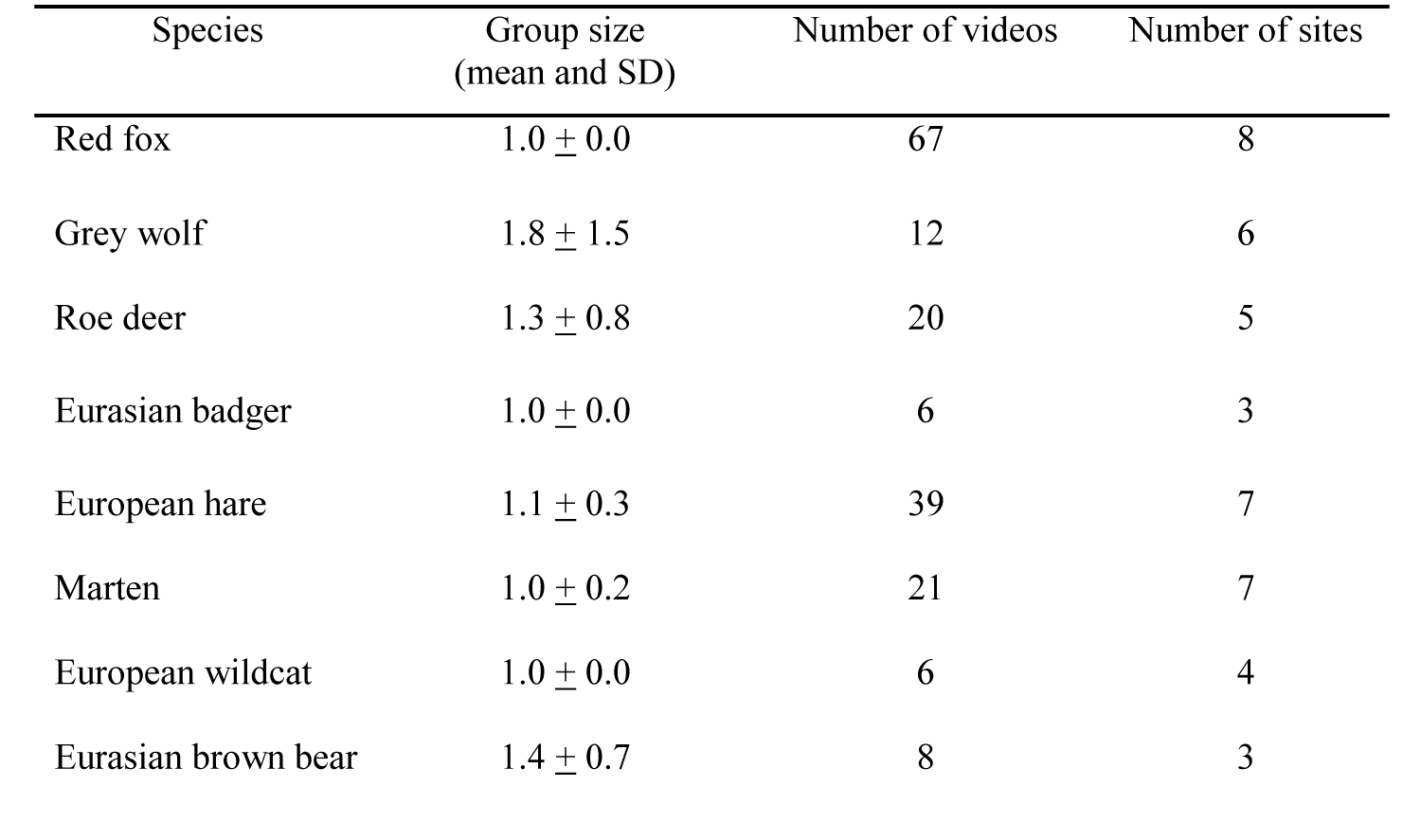
Estimates of average group size for each species derived from videos.

| Species | Group size<br>(mean and SD) | Number of videos | Number of sites |
| --- | --- | --- | --- |
| Red fox | 1.0 ± 0.0 | 67 | 8 |
| Grey wolf | 1.8 ± 1.5 | 12 | 6 |
| Roe deer | 1.3 ± 0.8 | 20 | 5 |
| Eurasian badger | 1.0 ± 0.0 | 6 | 3 |
| European hare | 1.1 ± 0.3 | 39 | 7 |
| Marten | 1.0 ± 0.2 | 21 | 7 |
| European wildcat | 1.0 ± 0.0 | 6 | 4 |
| Eurasian brown bear | 1.4 ± 0.7 | 8 | 3 |

The density at each study site (Table 4), and for all sites combined (Table 5), was calculated for all species except for golden jackal, as only two records were obtained at a single site. The density estimates for one site (Junik National park) were high for some species (fox, wild boar, marten and brown bear) compared to the other sites. The national brown bear population estimate in particular was considered to be high, largely due to the influence of the Junik data. When the Junik data were excluded from the analysis, a mean density of 0.12/km^2^ (range 0.04-0.20) and a mean population size of 488 (range 182-850) was calculated for brown bear.

**Table 4.** The mean density estimates (associated 95^th^ percentile ranges) calculated from the camera survey data and REM and the perceived frequency (frequent: F, sporadic: S or not present: NP) by land managers for each species at each site. Species recorded as present by the camera survey are shaded mid-grey and by the land manager survey as dark grey.

| Survey period | Site | Mean density, 95<sup>th</sup> percentile range (individuals/km<sup>2</sup>) and perceived frequency for each species |  |  |  |  |  |  |  |  |
| --- | --- | --- | --- | --- | --- | --- | --- | --- | --- | --- |
|  |  | Red fox | Grey wolf | Wild boar | Roe deer | Eurasian badger | European hare | Pine/beech marten | Wildcat | Eurasian brown bear |
| 1 | Malisheva | **0.17**<br>(>0.00-0.50) | **0.18**<br>(>0.00-0.44) | **0.00** | **0.00** | **0.00** | **0.69**<br>(>0.00-2.06) | **0.00** | **0.00** | **0.00** |
|  |  | F | F | F | F | NP | NP | NP | NP | NP |
|  | Shtime | **1.00**<br>(>0.00-2.27) | **0.23**<br>(>0.00-0.53) | **0.00** | **14.14**<br>(4.70-25.08) | **0.41**<br>(>0.00-0.83) | **4.12**<br>(> 0.00-10.02) | **0.83**<br>(> 0.00-1.66) | **0.43**<br>(>0.00-0.86) | **0.23**<br>(>0.00-0.68) |
|  |  | F | F | F | F | F | F | F | S | NP |
|  | Jezerc | **0.42**<br>(>0.00-0.99) | **0.08**<br>(>0.00-0.23) | **0.00** | **1.55**<br>(>0.00-4.70) | **0.00** | **0.58**<br>(>0.00-1.77) | **0.00** | **0.00** | **0.00** |
|  |  | F | F | F | F | F | F | F | S | S |
|  | Kamenica | **0.00** | **0.00** | **1.58**<br>(>0.00-4.77) | **0.00** | **0.00** | **0.55**<br>(>0.00-1.65) | **0.00** | **0.00** | **0.00** |
|  |  | F | F | F | F | F | F | NP | S | S |
|  | Pristina | **0.67**<br>(>0.00-1.49) | **0.09**<br>(>0.00-0.27) | **0.65**<br>(>0.00-1.99) | **0.00** | **0.00** | **0.00** | **0.20**<br>(>0.00-0.58) | **0.00** | **0.00** |
|  |  | F | F | F | F | F | NP | F | NP | NP |
|  | Dragash | **0.00** | **0.07**<br>(>0.00-0.21) | **0.53**<br>(>0.00-1.59) | **0.00** | **0.00** | **0.00** | **0.00** | **0.00** | **0.00** |
|  |  | F | F | F | F | NP | NP | F | S | F |
| 2 | Dubocak | **0.24**<br>(>0.00-0.58) | **0.00** | **0.76**<br>(>0.00-1.73) | **2.64**<br>(0.53-5.30) | **0.00** | **0.99**<br>(>0.00-2.59) | **0.06**<br>(>0.00-0.18) | **0.05**<br>(>0.00-0.15) | **0.46**<br>(0.15-0.84) |
|  |  | F | F | F | F | F | F | F | S | F |
|  | Junik | **4.14**<br>(1.28-7.45) | **0.11**<br>(0.03-0.23) | **6.19**<br>(2.77-10.00) | **4.73**<br>(1.18- 9.40) | **0.21**<br>(>0.00-0.57) | **0.00** | **1.24**<br>(0.26-2.63) | **0.22**<br>(>0.00-0.54) | **1.01**<br>(0.34-1.94) |
|  |  | F | F | F | F | F | F | F | F | F |
|  | Istog | **0.66**<br>(0.16-1.31) | **0.04**<br>(> 0.00-0.13) | **0.00** | **12.64**<br>(2.71-29.80) | **0.00** | **2.35**<br>(> 0.00-5.09) | **0.10**<br>(> 0.00-0.30) | **0.00** | **0.13**<br>(>0.00-0.39) |
|  |  | F | F | F | F | F | S | F | F | S |
|  | Mitrovica | **0.28**<br>(0.05-0.53) | **0.18**<br>(>0.00-0.47) | **0.19**<br>(>0.00-0.63) | **0.00** | **0.04**<br>(>0.00-0.15) | **2.76**<br>(0.21-3.24) | **0.16**<br>(>0.00-0.43) | **0.00** | **0.00** |
|  |  | F | F | F | S | S | S | F | F | S |
|  | Podujevo | **1.19**<br>(0.25-2.32) | **0.00** | **1.00**<br>(>0.00-2.25) | **0.00** | **0.12**<br>(>0.00-0.30) | **5.14**<br>(0.26-12.22) | **0.46**<br>(>0.00-1.32) | **0.13**<br>(>0.00-0.32) | **0.00** |
|  |  | F | F | F | F | F | S | F | F | S |

**Table 5.**
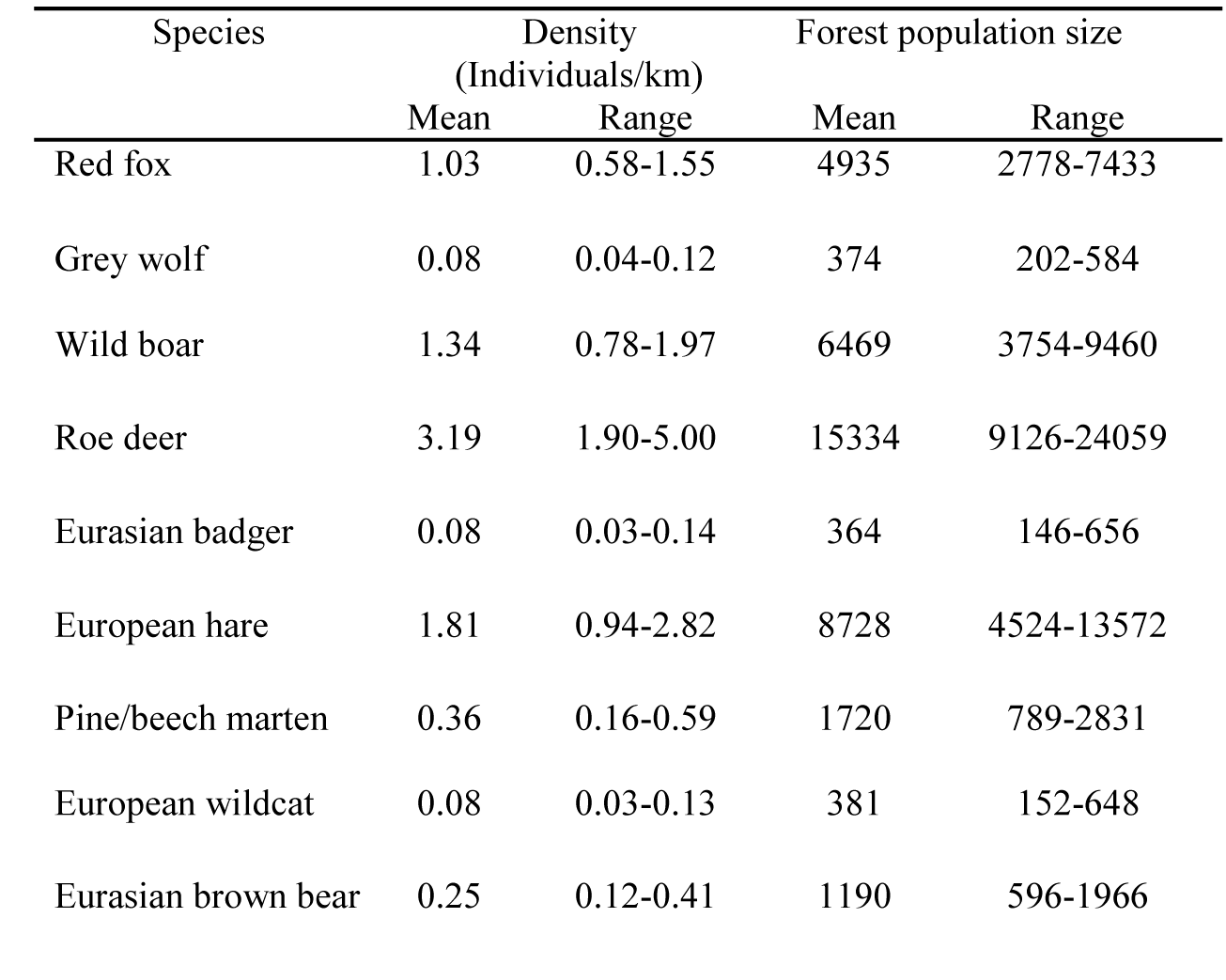
The mean densities, 95th percentile ranges and population estimates for mammal species in Kosovo forests calculated across all of sites and for all sites except Junik.

### Land manager surveys

The frequency of each species at each site, as assessed by the land manager’s survey, was compared to the data collected by the camera survey (Table 4). The red fox, grey wolf and wild boar were recorded as frequent at every study site, roe deer as frequent at ten sites and badgers and marten as frequent at eight and nine sites respectively. Wildcat and brown bear were assigned as either sporadic or not present at the majority of sites. All species were recorded by land managers as present at every site in survey period 2, while in survey period 1, on 12 occasions (5 species, 5 sites) a species was perceived as not present.

There were 17 occasions (8 species, 5 sites) in survey period 1, and 10 occasions (8 species, 4 sites) in survey period 2, where a species perceived as frequent by a land manager was not recorded by the camera survey. European hare and brown bear were recorded on one occasion each as not present by a land manager, but were recorded by the camera survey.

In addition to those recorded by the cameras, mammal species recorded at each site by land managers were, golden jackal (frequent in Istog and sporadic in Podujevo, Dubocak and Jezerc), chamois (*Rupicapra rupicapra*: frequent in Dragash and Junik), red deer (*Cervus elaphus*: sporadic in Shtime, Podujevo and Malisheva), lynx (*Lynx lynx*: sporadic in Dragash and Shtime) and rabbit (*Oryctolagus cunniculus*: sporadic in Pristina and Kamenica).

## Discussion

The main species of rabies concern, the red fox, was found by both the land manager survey and the camera trap survey to be the most widely spread medium to large wild mammal in Kosovo forests. It was identified by cameras at nine of eleven study sites and categorised as frequent by land managers at every site, while the camera survey provided additional information on relative densities between sites and therefore potentially different disease risk levels. The highest density was calculated at the most species rich site, the national park site of Junik, where hunting is permanently prohibited; while the lowest density was found at the least species rich site, Malisheva, which had one of the highest levels of public access and consisted of smaller, more fragmented areas of forest. The range of density estimates obtained from the camera survey for foxes in Kosovo forests are comparable to densities reported in other European countries, with rural fox densities ranging from 0.2 to 0.4/km^2^ in Northern Scandinavia [24] up to 4/km^2^ in South West England [25]. The total Kosovo fox population is likely to be much higher than the above estimate, as generally forest areas will support lower fox densities than more heterogeneous environments such as agricultural land [26].

In addition the camera survey provided information on specific locations where foxes and wolves co-existed at relatively high densities and/or in close proximity to human habitations, information that could inform disease management strategies. The grey wolf was the second most recorded species in Kosovo by the camera survey. It was recorded at eight sites throughout the country by the camera survey and was perceived as frequent at every study site by land managers. The mean density of grey wolf in Kosovo (0.08/km^2^) appears realistic and although high by European standards, for example 0.7-3.0/100 km^2^ has been recorded in Poland [27] was not exceptional when compared with the densities of 0.02-0.10/km^2^, calculated from cull numbers in the Social Federal Republic of Yugoslavia in 2000 [28]. The estimated forest grey wolf population, 374 (range 20–584), can be considered to be a national estimate, as wolves are unlikely to exist in substantial numbers outside of the surveyed habitat. This is high compared with the estimate for Kosovo in 2003 of up to 100 individuals [9] and the figure of 800 individuals reported for Serbia (which is eight times the size of Kosovo) in 2011 [29].

It is likely that the Kosovo grey wolf population is increasing. The European grey wolf population quadrupled in size between 1970 and 2005 [30] and the Dinaric-Balkan wolf population, with an estimated 3,900 individuals, is one of the two largest grey wolf populations on the continent, crossing the most national borders [29]. The grey wolf has also become an important conservation species and is now protected in Europe under various legislation [30] and there has also recently been an increase in legislation aimed at protecting biodiversity in Kosovo [31]. Additionally, the roe deer, one of the grey wolf’s main prey species, was found by the camera survey to be the most abundant medium to large mammal species in Kosovo and had a mean density of (3.5/km^2^), that was low compared to estimates recorded for other European countries, for example 7/km^2^ in the Czech Republic [32], while the mean population estimate of 15,000 is high when compared with the Kosovo Government estimate of 5000-6000 roe deer for Kosovo in 2003 [9]. The discrepancy between these two results could be explained by the fact that there has the Kosovo Government has introduced initiatives such as hunting bans to moderate the impact of illegal hunting.

Roe deer were perceived as frequent by land managers at 10 out of the 11 sites surveyed but only found present at 5 sites by the camera surveys. There were further inconsistencies between the land manager surveys and camera survey for mustelids; the land manager survey recorded badgers and marten as frequent at eight and nine sites respectively, while the camera survey recorded badgers at four sites and marten at seven. This may have been due to the fact that land managers surveys covered a much greater time period than the camera survey or could suggest that further effort was required for the camera survey. The discrepancies are fewer in survey period 2 when the number of camera days was increased. The estimated mean badger density 0.8/km^2^ however appears realistic as it is similar to badger densities found in Poland (0.04-0.10/km^2^) and the Czech republic (0.05-0.12/km^2^) [33]. The mean density estimate for marten (0.34/km^2^) also appears realistic as it is similar to pine marten densities of 0.34/km^2^, calculated in Italy also using the REM [34] and 0.54 km^2^ recorded in the Białowieża forest, Poland [35].

Wild boar can host diseases such as CSF and ASF and are also an important game species. Land managers reported wild boar as frequent at every site and they were recorded by the camera survey at seven sites. The mean density estimated for Kosovo (1.34/km^2^) is similar to densities recorded in Russia (1.2-1.9/km^2^), Italy (1.4-1.7/km^2^) and France (1.0-2.9/km^2^) [36] and the lower of the threshold density ranges (0.6 and 1.1/km^2^) reported for CSF persistence [37]. By far the highest density (6.19/km^2^) was recorded at the National Park site of Junik while there was an absence of boar records at the three sites with greater public access. The mean forest population of wild boar was estimated at 6469 individuals, which appears realistic when compared with an estimate of 30,000 wild boar in Serbia in 2004/2005 [30] and an estimate of 6000-10,000 for Kosovo in 2003 [9].

As well as species of disease concern, the surveys also provided data on species of conservation concern. The International Union for the Conservation of Nature (IUCN) has designated status for brown bear in the Balkan region as ‘vulnerable’ [30]. The brown bear was classed as present by land managers at eight sites and by camera surveys at four sites. The brown bear was the only species for which the mean density (0.25/km^2^) and mean population estimate (1190) were the considered unrealistic. In the last five years the bear population in Serbia has been estimated at approximately 80 individuals [29] and in 2003 the Kosovo bear population was recorded as between 80 and 100 [9], though the accuracy of both of these values is unknown. The site with the highest density of bears, the National Park site of Junik, was not considered proportionately representative of bear suitable habitat in Kosovo. When Junik was removed from the data, the mean population estimate was 488, which appears high but more realistic. Brown bear were the species recorded at the lowest maximum distance from the cameras (three metres). There is therefore some evidence bears may have been attracted to the cameras thus increasing their number of records. It is also possible that the DDM used for brown bear (6 km) was too low for Kosovo. There are limited published data on the range sizes of European brown bear. These finding suggest that the camera survey method may require further refinements to make it suitable for the assessment of some species.

The European hare has been identified as an important conservation species due to its decline across Europe since the 1960s [38]. In this study, hare densities were found to be low across the eight sites at which they were recorded at, though forest is not an optimum habitat for hares [39]. The fact that hare were recorded at a high proportion of sites would suggest the total Kosovo hare population is likely to be high, and in excess of the mean forest population estimate of 8728. Another important conservation species is the European wildcat. Most European populations analysed have showed evidence of hybridisation with the domestic cat *Felis catus* and it is estimated that 88% of wildcats in Scotland may be hybrids [40]. Wildcats were recorded by the camera surveys at four sites and were recorded as present at nine sites by land managers. The difficulty of identifying true wildcats from hybrids makes estimates for this species uncertain. The Kosovo wildcat mean density was calculated as 0.08/km (range 0.03 to 0.13), which is low compared to Sicily (0.22 to 0.44/km2):, but similar to Poland [0.1 to 0.13/km2: 41].

Two records of Golden jackal were obtained from the site of Podujevo near the north eastern Kosovo/Serbian border. This is the first confirmed live record of golden jackal in Kosovo. There was estimated to be between 4500 and 5000 golden jackals in Serbia in 2011 [30] and densities up to 1 group/km^2^ have been recorded in the Balkans [42]. This is a significant finding however, as golden jackals are currently expanding their range across Europe and recently the first evidence of their recolonisation of FYROM, since their extinction in the 1960s, was obtained through camera traps [43].

Overall the density estimates produced by the REM for each species, with the exception of the brown bear, appear realistic and are comparable to those published for similar environments in other European countries. Unlike previous methods used in Kosovo, the estimates provide a standardised measure of relative abundance for each species at different sites across the country. Despite the improvements made to the data collection methodology in survey period 2, the 95^th^ percentile ranges calculated are wide and the methods would require further refinement if the results were to be used to look at the demographics of wild mammal species in more detail. For this study, it was necessary to make a number of modifications to the REM in order to adapt it to the time and resources available and to make the methods practicable for local stakeholders. Sampling effort will affect the accuracy and precision of most population estimation techniques. For robust estimates using the REM, a minimum of 20 cameras and a minimum of 10 records per species is recommended [12]. This was achieved for the combined Kosovo forest data, but not for individual sites. A study of Harvey’s duiker [44] suggested the minimum number of camera days for satisfactory precision was 250 to 300 (for densities between 12 and 15/km^2^). For the second survey period, the number of camera days was increased to an average of 350 days per site, however for the density levels estimated in this study; a greater amount of sampling effort would be required for increase precision.

Increased sampling effort would also decrease the effect of other potential sources of bias. Due to time and resource limitations, the camera surveys were conducted using transects of cameras, rather than grids, over short periods of time and at different times of year for different sites. At some of the sites assessed in the first survey period, the cameras were placed much further apart than the 330 m distance recommended [12]. The detection probability of some species may have therefore been reduced as a result. The land manager surveys, particularly in survey period 1, found some species present at some sites that were not recorded by the cameras. However, the land manager surveys also indicated that there were overall fewer species present at the sites surveyed in period 1 than period 2. The detection probability of some species at some sites may also have been reduced by temporal factors including seasonality, even though young juveniles were not included in the analysis. Ideally, given adequate resources, surveys should all have been conducted in spring or autumn, when mammal species are at their most active and detection probability is at its highest. In contrast, the longer time period and geographic scale over which land managers recalled sightings when surveyed is likely to have biased their perception of species frequencies upwards. This is a source of bias that is likely to affect species data that are derived from observations. One or a combination of these factors could explain why land manager surveys reported the presence of species such as chamois that were not identified by the cameras.

Species specific factors such as body size and animal behaviour are important sources of bias in detection probability, particularly for population estimation methods such as direct counts. In this study, the same methodology was used for a variety of species and the camera radius and angle of detection were calculated retrospectively using methods that were not species or camera location specific, which may have biased the REM estimates. There was evidence from the photograph analysis that the behaviour of some species e.g. the apparent attraction of brown bears towards the cameras, may have affected their detectability and that site factors such as vegetation cover also influenced animal detectability. Both of these factors were found to be more influential for species detection probability than mammal body size. Recently, novel methods for calculating camera detection probability have been developed, that aim to reduce bias in REM estimates, by accounting for species specific influences such as body size and camera location specific influences such as vegetation cover [45]. It is recommended that these methods are employed, if practicably possible, in future camera surveys to improve REM estimate accuracy.

Similarly, obtaining accurate values for the DDM and group size for each species was difficult as few data of this kind were available for mammals in Kosovo. For movement data, websites such as Movebank and collaborations such as Euroungulates are invaluable, as they provide access to large amounts of long-term data for various species in a range of habitats. It is unknown how representative the radio-telemetry or GPS daily movement values used were to Kosovo species and it is acknowledged that interval locations will underestimate daily movement considerably and particularly for smaller species of mammals [12] and thus overestimate density. The mean DDM values obtained in this study in fact varied very little between most species, despite the wide variety of data sources, however for species with large, site specific and variable home ranges, such as the grey wolf, the degree of associated error will be much greater. There are however currently no viable alternative methods for estimating DDM for such species, but any new DDM estimates could be applied to the data to revise the above density estimates.

In this study, using videos to estimate group size provided a greater degree of site specificity, compared to using values from the research literature. For species with larger group sizes, it is recognised that it is more likely that individuals will be missed by the cameras, therefore biasing estimates low. It was for this reason that the wild boar group size for this study was taken from the research literature and could therefore be a potential source of bias. The other species in the study, with the exception of the grey wolf, generally moved around singularly or in small groups, therefore the associated bias is likely to be minimal. Due to the variability and potentially large sizes of grey wolf groups, it is recognised that this estimate is likely to have been biased low as a result. Ideally, a site specific, independent measure of group size is required to improve estimate accuracy, for example, a number of cameras could be deployed in one location positioned so as to obtain several estimates of numbers of individuals for the same group of animals. Overall it is recommended that further research into the influence of species specific factors on estimate bias is required.

The population estimates for Kosovo forests appear high for some species when compared with previous estimates in the region from hunting bags and direct counts; however the accuracy and precision of these methods are often unmeasurable and they are subject to a number of different sources of bias. It is possible that the high estimates for some species in Kosovo may be symptomatic of a general increase in wildlife density across Europe [46]. Camera placement should be random and representative of the total area studied. Particularly in the second survey period of the study, guidance was provided to local stakeholders to ensure that the location of each camera transect was representative of the wider study area in which they were placed. This was not always possible where sites were difficult to access and it is recognised that, given more resources, a grid of cameras would have provided a more random representation of each study area. Overall species detection increased in the second survey period, so there is little evidence that camera placement biased the population estimates.

Despite the provision of methods and training, a degree of operator bias is likely to have occurred in this study, with more experienced operatives potentially placing cameras in positions that procured more data. It is recognised that citizen science initiatives are susceptible to such biases, related to the level of skill, experience and dedication of volunteers [47] however these kinds of bias are likely to be even greater for population estimation techniques that rely on land manager surveys, hunting bags and direct counts; for example in this study there was some evidence that land managers were more likely to perceive species they hunt such as wild boar and fox as frequent, compared with those they didn’t, such as badger and wildcat and that species they had little interest in could potentially be misidentified, e.g. hare for rabbit.

Overall this study found that camera surveys using local stakeholders can provide data more applicable to disease and conservation management, when compared with traditional methods of estimating wild mammal species presence and abundance. Camera surveys are less susceptible to biases, particularly those related to human operators, and this is one of the most suitable methods to estimate density [48]. Guidance on estimation of wild b. However, this method still requires further refinement and it is recommended that for more robust camera survey population estimates the have a greater degree of utilisation, there should be further research into the measurement and moderation of different sources of bias and a greater number and variety of sites should be sampled. If these points were addressed and the camera survey was extended to other habitat types, robust national estimates for all medium to large wild mammal species in Kosovo would be attainable.

## Conclusion

The combined use of land manager surveys and camera surveys, delivered via local stakeholders, together with the REM has provided the first systematic distributions, densities and population estimates for wild mammal species in Kosovo. This includes the first live record of golden jackal. Although sources of bias have been identified, these estimates are likely to be more accurate than methods such as direct counts and hunting bag estimates. It is acknowledged that precision and accuracy could be further improved through increased sampling effort and method refinement to address likely sources of bias, nonetheless the findings of this study suggest that, with a moderate amount of development, camera trapping can be used as an effective and practicable method to estimate national distributions and population sizes of medium to large sized wild mammals.

## Acknowledgements

The work described in this paper was undertaken through an EU funded project managed by the European Union Office in Kosovo, Project Number EuropeAid/127852/D/SER/KOS. The authors acknowledge the support received from the European Commission and would also like to thank Dr. Francesca Cagnacci (Edmund Mach Foundation) and Johannes De Groeve (Ghent University) from the Euroungulate network, Wiebke Ullmann (University of Potsdam and BioMove) and Professor Melvin Sunquist (University of Florida) for providing animal movement data for the analysis. The authors would also like to thank Bajram Batusha (Ministry of Agriculture, Forestry and Rural Development, Kosovo), the members of the Kosovo Forestry Agency, Kosovo hunting associations, managers of licensed hunting grounds and National Parks, US members of the Kosovo Force (KFOR), who facilitated the collection of field data and project delivery.

